# The h-current controls cortical recurrent network activity through modulation of dendrosomatic communication

**DOI:** 10.1101/2023.07.12.548753

**Authors:** Yousheng Shu, Andrea Hasenstaub, David A. McCormick

**Affiliations:** The Fudan University Fenglin Campus, 131 Dong’an Road, Xuhui District, Shanghai; Department of Otolaryngology-Head and Neck Surgery (OHNS), University of California, San Francisco, 675 Nelson Rising Lane, #514B, San Francisco CA 94158; Department of Neuroscience, Yale University School of Medicine, New Haven, CT 06510; Institute of Neuroscience, University of Oregon, Eugene, OR 97403

**Keywords:** Persistent Activity, Neocortex, Dendrosomatic, h-current, Memory, Gain

## Abstract

A fundamental feature of the cerebral cortex is the ability to rapidly turn on and off maintained activity within ensembles of neurons through recurrent excitation balanced by inhibition. Here we demonstrate that reduction of the h-current, which is especially prominent in pyramidal cell dendrites, strongly increases the ability of local cortical networks to generate maintained recurrent activity. Reduction of the h-current resulted in hyperpolarization and increase in input resistance of both the somata and apical dendrites of layer 5 pyramidal cells, while strongly increasing the dendrosomatic transfer of low (<20 Hz) frequencies, causing an increased responsiveness to dynamic clamp-induced recurrent network-like activity injected into the dendrites and substantially increasing the duration of spontaneous Up states. We propose that modulation of the h-current may strongly control the ability of cortical networks to generate recurrent persistent activity and the formation and dissolution of neuronal ensembles.

## Introduction

A characteristic feature of the cerebral cortex is the presence of strong recurrent excitatory and inhibitory connections, with each cortical pyramidal cell receiving tens of thousands of synaptic inputs from many thousands of other cortical neurons (Abeles, 1991; White, 2012; Braitenberg and Schüz, 2013). This massive recurrent network connectivity provides to each neuron the synaptic context under which the cell operates and allows for a nearly infinite variety of neuronal associations that form co-active and interacting ensembles of cortical neurons (reviewed in (Haider and McCormick, 2009)).

A prominent feature of this recurrent connectivity within the cortex is the generation of persistent activity. One example of persistent activity is observed during slow wave sleep and anesthesia, in which cortical networks generate periods of recurrent activity separated by relative silence, the so-called Up and Down states (Metherate and Ashe, 1993; Steriade et al., 1993, 2001; Cowan and Wilson, 1994). The Up state is generated by activation of cortical recurrent networks, which have a tendency, owing to their massive recurrent connectivity, to rapidly transit from the Down to Up states. During the Up state, the bombardment of each cortical pyramidal cell with EPSPs is rapidly balanced by adjustments in the activation of IPSPs, resulting in a semi-stable state within the cortical neuron and network (Shu et al., 2003a, 2003b).

Another example of persistent activity is observed during active movement and arousal in mice, in which cortical neurons exhibit prolonged depolarization mediated by barrages of synaptic potentials arising from activity in corticocortical and thalamocortical networks. These barrages of synaptic activity can strongly influence sensory evoked responses and correlated with changes in the ability to perform sensory detection/discrimination tasks (Sachidhanandam et al., 2013; McGinley et al., 2015; Neske et al., 2019; McCormick et al., 2020; Nestvogel and McCormick, 2022). In higher mammals, such as cats, persistent activity associated with barrages of synaptic activity in the waking state are also observed (Steriade et al., 2001), and similar activity was observed primate V1 neurons upon visual stimulation (Tan et al., 2014).

The ability to generate persistent activity in cortical and thalamocortical networks has been proposed to be a critical property in the operation of the cortex - allowing, for example, the generation of working memory, planned movements, and context-dependent gain modulation (Major and Tank, 2004; Wang et al., 2007; Fuster, 2021; Khona and Fiete, 2022). The ability to generate persistent activity within the cortical and thalamocortical network depends on the interactions of not only neurons, but also the subcomponents of those neurons, such as dendritic-somatic-axonal communication (Bernander et al., 1991; Destexhe et al., 2003; Shu et al., 2003a; Major and Tank, 2004; Wolfart et al., 2005).

A wide variety of factors may influence the generation of recurrent network activity, including the activation of conductances intrinsic to the participating neurons (Egorov et al., 2002; Compte et al., 2003; Major and Tank, 2004) or changes in synaptic strength between these cells (reviewed in (Abbott and Regehr, 2004)). One potentially important factor is the ability of synaptic inputs arriving in the dendrites to influence the generation of action potentials, which, in adult cortical pyramidal cells, are typically initiated in the axon or axon initial segment/somatic compartment (Turner et al., 1991; Stuart et al., 1997a, 1997b; Shu et al., 2007; Foust et al., 2010; Popovic et al., 2011).

Immunocytochemical localization HCN1 and HCN2 subunits of the h-channel in the cerebral cortex reveal a distinct pattern (Lörincz et al., 2002; Notomi and Shigemoto, 2004). The apical dendrites of layer 5 pyramidal neurons exhibit a progressive increase in HCN1/2 channel density as the dendrite approaches layer 1, suggesting that the h-current may be particularly prominent in distal dendrites. This hypothesis was confirmed with apical dendritic patch clamp recording (Magee, 1998; Berger et al., 2001), which also demonstrated that the presence of a large h-current in distal dendrites significantly decreased the temporal integration at these locations (Schwindt and Crill, 1997; Magee, 1998, 1999; Stuart and Spruston, 1998; Williams and Stuart, 2000, 2003; Berger et al., 2001, 2003; Oviedo and Reyes, 2005). These results indicate that the h-current has a powerful effect on dendrosomatic integration and communication (Magee, 1998, 1999; Poolos et al., 2002; Nolan et al., 2004; George et al., 2009; Harnett et al., 2015; Kelley et al., 2021; Routh et al., 2022). Here we tested the functional consequences of this hypothesis by examining the effects of alteration of the h-current on the generation of the recurrent network activity of the Up states in vitro, since this activity depends critically upon balanced recurrent excitation and inhibition in local cortical networks (Sanchez-Vives and McCormick, 2000; Shu et al., 2003a, 2003b).

## Experimental Procedures

### Slice Preparation

Oblique slices from somatosensory cortex or coronal slices from prefrontal cortex were prepared from 7-12 week old ferrets. Ferrets were initially deeply anesthetized with pentobarbital (30 mg/kg) and decapitated, and the brain was quickly exposed and removed from the connected tissue in ice-cold oxygenated (95% O_2_ and 5% CO_2_) sucrose-substituted artificial cerebrospinal fluid (ACSF) in which sucrose was used as a substitute for NaCl. Slices for use in the interface chamber were cut (400 μm thick) and transferred to the chamber; they were then perfused with normal and sucrose-substituted ACSF (1:1) for 20 minutes at 36 ºC, then incubated in normal ACSF for 1 hour. After incubation, slice solution was modified to contain 1 mM MgSO_4_, 1 mM CaCl_2_ and 3.5 mM KCl before recording (modified ACSF). For recordings in the submerged chamber, slices were cut (300 μm thick) and immediately transferred to an incubation beaker filled with aerated normal ACSF containing (in mM): NaCl 126, KCl 2.5, MgSO_4_ 2, CaCl_2_, 2, NaHCO_3_ 26, NaH_2_PO_4_ 1.25, dextrose 10 (315 mOsm, pH 7.4) and held at 35 ºC until use. After at least 1 hour of incubation, slices maintained in the beaker were transferred to a submerged chamber containing aerated modified ACSF; both a bottom grid and a top grid were used to lift and hold the slices in position, and cortical neurons were visualized with an upright infrared-differential interference contrast (IR-DIC) microscope (BX61WI, Olympus). A light sensitive camera (OLY-150, Olympus) was used for tracing the fluorescent neuronal profiles.

All experiments were done in modified ACSF at a temperature of 36-36.5 ºC; most of the slices maintained in this solution generated spontaneous recurrent network activity.

### Electrophysiological recordings

In submerged slices, whole-cell recordings were achieved from both soma and apical dendrites using a Multiclamp 700B or Axoclamp 2B amplifier (Axon Instruments, Union City, CA). Patch pipettes were formed on a P-97 microelectrode puller (Sutter Instruments, Novato, CA) from borosilicate glass (1B200-4, WPI, Sarasota, FL). Pipettes for somatic recording had an impedance of 5-6 MΩ, and were filled with an intracellular solution that contained (in mM): KGluconate 140, KCl 3, MgCl_2_ 2, Na_2_ATP 2, HEPES 10, EGTA 0.2, pH 7.2 with KOH (288 mOsm). Alexa Fluo 488 (100 μM) and biocytin (0.2%) were added to the pipette solution for tracing and labeling the recorded cells. Approximately 10 minutes after whole-cell recording was established, the structure of the dendritic tree was examined under the fluorescent microscope equipped with a 60x water objective and a magnifier up to 2×. Only those pyramidal neurons with most of the dendritic tufts preserved were used in our study. Patch pipettes for dendritic recording were filled with a similar intracellular solution, but without fluorescent dye added; these electrodes had an impedance of 9-15 MΩ. The pipette was advanced to the selected dendritic segment with a positive pressure of about 110 mbar, and guided by switching back and forth between the fluorescent and DIC images of the dendrite. The dendritic segment was then pressed by the pipette tip and a negative pressure was applied to form a seal. As soon as a tight seal (>10 GΩ) was achieved, pulses of brief suction were applied to break the membrane, and whole-cell configuration could be easily obtained. Thereafter, steps of positive current and negative current were injected to the soma and dendrite to examine the intrinsic membrane properties of the recorded neuron. During the whole period of simultaneous somatic and dendritic recordings, access resistance was monitored frequently; recordings with access resistance higher than 25 MΩ for somatic recording, 45 MΩ for dendritic recording, were discarded. Bridge balance and capacitance neutralization were carefully adjusted before and after every experimental protocol.

In slices maintained in an interface chamber, extracellular multiple-unit recordings were obtained with a single tungsten electrode (300-500 KΩ, Fredereck Haer Corporation); intracellular recordings were carried out with beveled glass microelectrodes with an impedance of 60-90 MΩ filled with 2 M potassium acetate. Both extracellular and intracellular recording electrodes were inserted in layer 5 and within 100 μm of each other. Intracellular signals were amplified with an Axoclamp-2B amplifier. Again, bridge balance and capacitance compensation were monitored continuously and adjusted as required. In all experimental conditions, intracellular recordings were accepted if they showed a stable membrane potential below -55 mV at rest and exhibited the ability to generate a train of action potentials upon depolarization. In both sharp electrode and whole cell patch clamp recordings, only data collected from regular spiking neurons were included for analysis. The recorded signals were acquired, digitized, and analyzed with Spike2 system and software (CED, Cambridge, UK). The h-channel blocker, ZD 7288, was applied through bath perfusion at a low concentration of 25 μM, to minimize the side effects such as its impact on synaptic transmission.

## Mathematical Methods

All computations were performed in Spike2 (Cambridge Electronic Design, Cambridge, UK) and MATLAB (MathWorks, Bethesda, MD). Power spectra were calculated using Welch’s method. Continuous frequency transfer functions were estimated using Welch’s averaged periodogram method. Discrete frequency transfer functions were estimated using Wiener’s method. All error bars were calculated as the standard error of the mean and data in the text are reported as mean +/-standard deviation.

A number of experiments were performed with a dynamic clamp technique using a DAP-5216a board (Microstar Laboratory). Noisy conductances were constructed according to an Ornstein-Uhlenhbeck (colored noise) model. For measurement of subthreshold transfer function, a conductance with reversal potential of 0 mV, a time constant of 5 ms, and whose standard deviation was approximately half its mean, was injected into the cell. The standard deviation (and, concurrently, the amplitude) of the conductance was adjusted to give a 10 mV peak-to-peak membrane potential deviation, and the holding current was adjusted to keep the cell just below spike threshold.

Transfer function from the injected conductance to the recorded membrane potential fluctuations was then calculated.

## Results

Simultaneous intracellular (regular spiking cells) and extracellular multiple unit recordings from layer 5 of the ferret prefrontal and somatosensory cortical slice maintained in the interface chamber revealed spontaneous Up and Down states, as reported previously (Sanchez-Vives and McCormick, 2000; Shu et al., 2003a, 2003b). Bath application of the h-current antagonist ZD7288 (20; n=3, 50; n=7,or 100 μM; n=5) resulted in a nearly 500% increase in duration of the Up states from an average of 0.84 (+/-0.25; S.D.) to 4.1 (+/-1.3) sec, and a smaller 63% increase in duration of the Down state from 6.5 +/-1.6 sec to 10.6 +/-4.4 sec, in both the intracellular and extracellular multiple unit recordings, over a period of 20-60 minutes (see Supplement 1). Given the prominent localization of HCN1/2 h-current subunits in the apical dendrites of layer 5 pyramidal cells (Lörincz et al., 2002; Notomi and Shigemoto, 2004), we hypothesized that this large effect of ZD7288 may have resulted from actions on the dendrosomatic integrative properties of these neurons. Therefore to address this question, we developed a submerged slice preparation of the ferret somatosensory cortex that generates the slow oscillation (Figure 1), and that would allow the simultaneous recording of both somatic and apical dendritic compartments from the same neurons.

**Figure 1.**
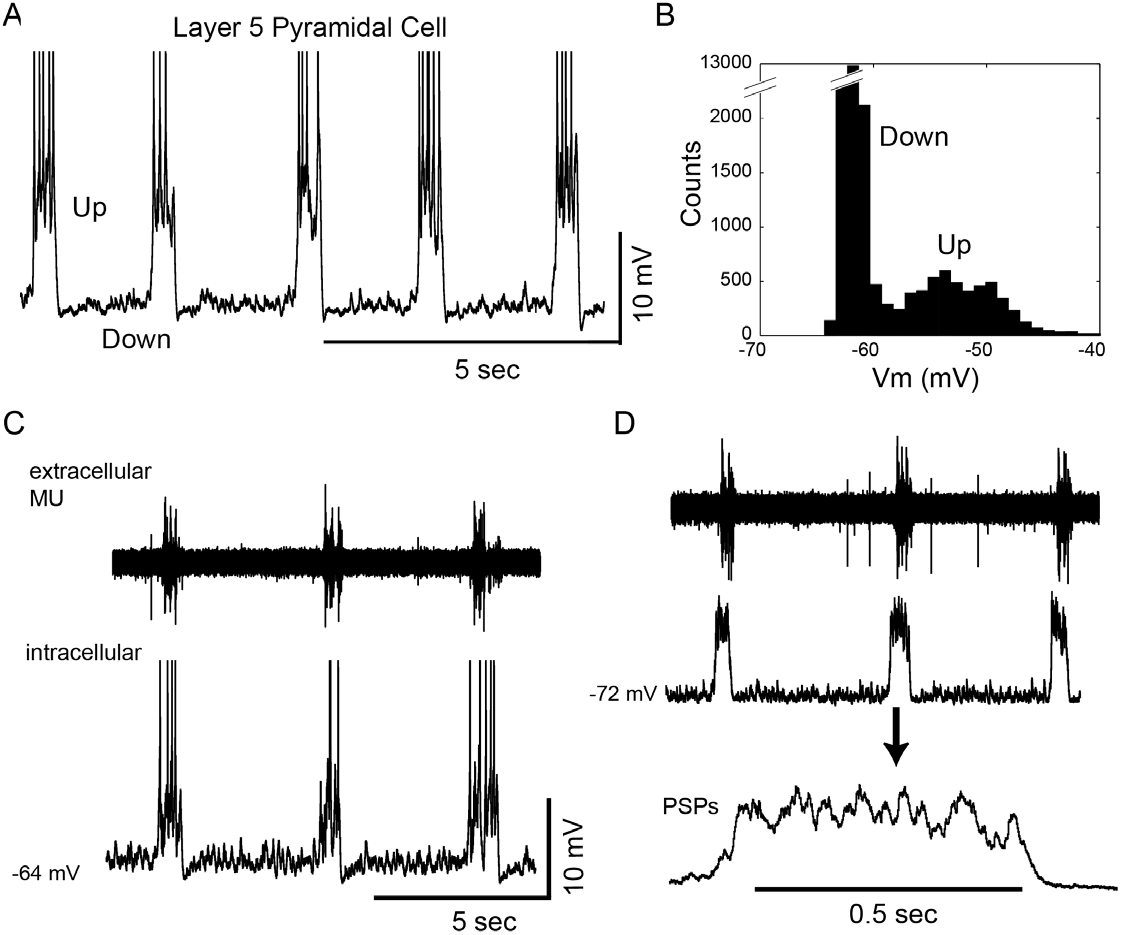
Characteristics of the slow oscillation in the submerged ferret somatosensory cortical slice. A. Slow oscillation recorded in a layer 5 pyramidal cell in a somatosensory cortical slice maintained in a submerged chamber. B. Histogram of the membrane potential illustrating the Up and Down states. C. Simultaneous extracellular and whole cell recording (pyramidal cell) from layer 5 in a submerged chamber demonstrating the rhythmic recurrence of Up states. D. Hyperpolarization of the pyramidal cell illustrates the barrages of PSPs underlying the Up state.

As in the interface chamber, whole cell recordings from layer 5 pyramidal cells in the submerged chamber revealed the spontaneous occurrence of Up and Down states that were characterized by the arrival of barrages of synaptic potentials during the Up state (Figure 1A, D; n=55). Simultaneous extracellular multiple unit and whole cell recording revealed that the Up states occur simultaneously with activity in many nearby neurons (e.g. see Figure 1C,D; n=5). During the Up state, pyramidal neurons received a 3-13 mV depolarizing barrage of synaptic potentials (Figure 1A,C), that resulted in action potential discharge of approximately 0-10 Hz. In comparison to the Up and Down states recorded in prefrontal cortical slices in interface chambers, the Up states in somatosensory cortical slices maintained in submerged chambers were typically less frequent, smaller in amplitude, and shorter in duration.

Simultaneous apical dendritic (80-600 μm from the soma) and somatic patch clamp recordings (n=50) from regular spiking layer 5 pyramidal neurons demonstrated that the Up state was associated with barrages of synaptic potentials that were visible in both somatic and apical dendritic compartments (Figure 2). Comparing the apical dendritic and somatic recordings revealed that the depolarizations associated with the synaptic barrages were similar in amplitude-time course, although the overall amplitude (as measured by the area under the synaptic barrage) was always (n=35/35 cells) larger at the somatic recording site (Figure 2B, C), especially in comparison to more distal apical dendritic locations (Figure 2D). The ratio of the area (mV.ms) under the Up state, between the somatic and apical dendritic compartments, decreased with increasing distance from the soma, and this decrease could be fit by an exponential function (λ = 980 μm; Figure 2D). Even though the amplitude and time course of the somatic and dendritic recordings were very similar, they were not merely scaled versions of one another. Subtraction of the dendritic recording from the somatic recording revealed small (< 3 mV) variations in membrane potential at the two recording sites (Figure 2B).

**Figure 2.**
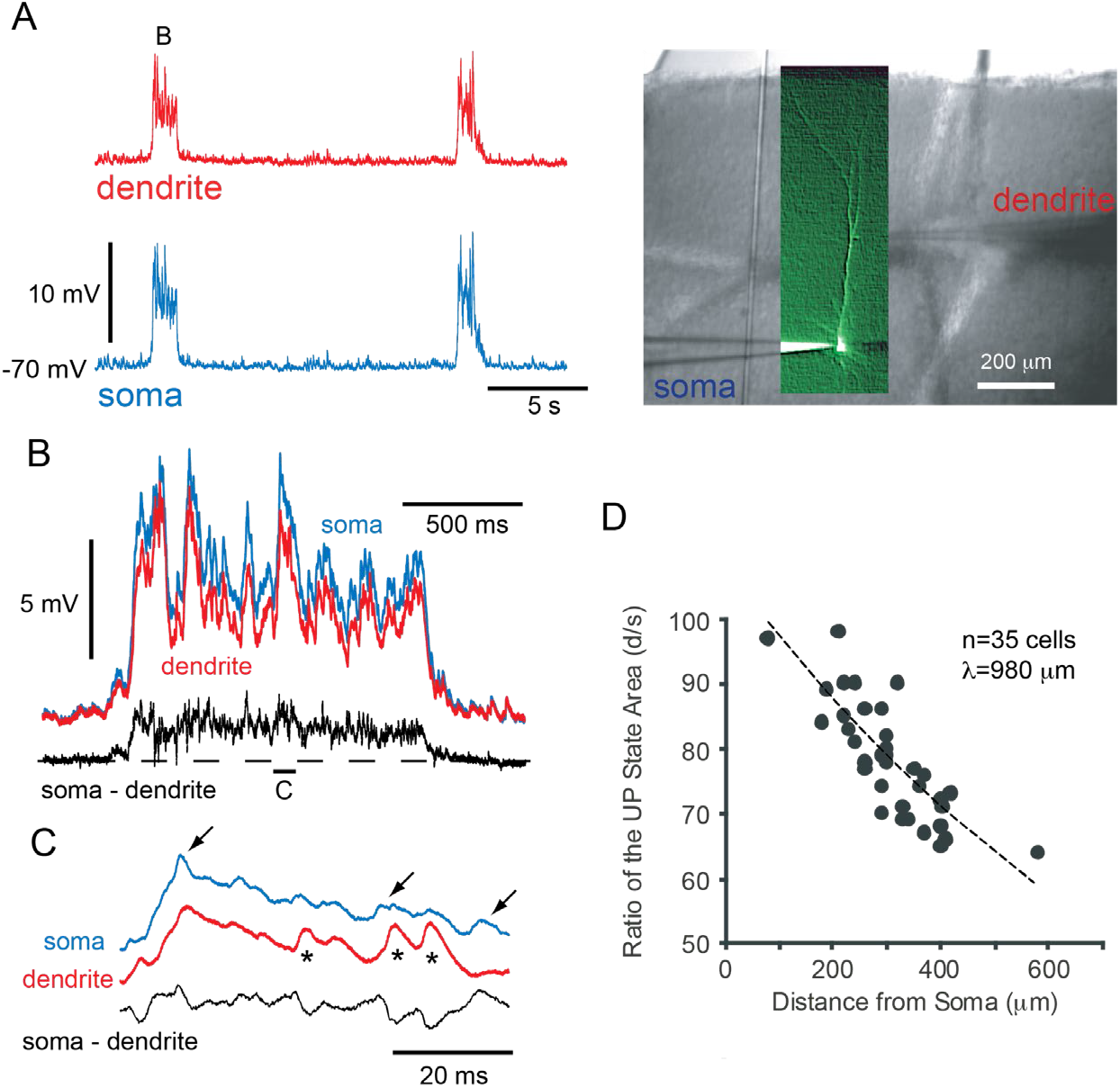
The synaptic activity of the slow oscillation is larger in the soma than that in the apical dendrite in vitro. A. Simultaneous somatic and apical dendritic recording from a large layer 5 pyramidal cell during the generation of the slow oscillation. The cell was lightly hyperpolarized with the injection of current to prevent the generation of action potentials. The dendritic recording was 285 μm from the soma. On the right is a combined fluorescent and DIC image of the recorded cell with the somatic and dendritic patch electrodes. B. Overlay of one Up state recorded simultaneously from the apical dendrite and soma. Subtraction of the dendritic recording from the somatic one yields a varying difference, indicating that they are not merely scaled copies of each other. C. Expansion of a portion of the Up state (as indicated in B). Arrows indicate events that are more distinct in the soma than the apical dendrite, while asterisks indicate events that are more prominent in the apical dendrite than in the soma. D. Ratio of the average area (mV.ms) under synaptic barrages of the Up state in the apical dendrite over the soma. Note that the somatic Up states are always larger and that the ratio decreases with distance from the soma in a manner that can be fit with an exponential function (λ of 980 μm).

Closer examination of these small variations revealed brief (<20 msec) events that were sometimes larger in the soma than in the apical dendrite, or vice versa (Figure 2C).

Some of these events resembled single PSPs in their shape, while others were more consistent with generation by compound PSPs (not shown). This result is consistent with the UP state being generated by synaptic barrages arriving in various locations on the cell, with the exact spatial-temporal pattern varying over the time course of the Up state (see also (Waters and Helmchen, 2004)).

In the submerged chamber, the bath application of ZD7288 (25 μM) during either extracellular multiple unit (Figure 3A; n=5) or whole cell recording from either the somatic or combined somatic/apical dendritic (Figure 3B; n=55) compartments of layer 5 pyramidal cells initially resulted in the reduction in rate, or temporary cessation, of the slow oscillation (although this effect was much less marked in the interface chamber, where the slow oscillation is more robust). Whole cell (submerged chamber) and intracellular (interface chamber) recordings revealed that this ZD7288-induced reduction in the slow oscillation occurred in conjunction with a progressive hyperpolarization of layer 5 pyramidal neurons (Figure 3B), to a final level of 3-14 mV (average of –9.3 +/-3.5 mV, n=5, interface; -9.1 +/-3.2 mV, n=7, submerged) below baseline. After 2-7 minutes, and in many cells while the hyperpolarization was still progressing, the Up state reappeared or increased its rate of occurrence. Following full wash-in of ZD7288, the Up state duration was markedly prolonged from a pre-ZD7288 average of 0.68 +/-0.3 sec to 2.3 +/-0.3 sec (submerged chamber; Figure 3A,C). The down state duration also increased from an average of 7.8 +/-3.6 to 11.5 +/-3.3 seconds (n=10), as had been observed in the recordings obtained in the interface chamber (see above). The hyperpolarization of cortical cells and increase in duration of the slow oscillation were observed even with concentrations of ZD7288 as low as 1-5 μM (n=3; submerged chamber; see Supplement-2). In addition to increasing the duration of the Up state, bath application of ZD7288 also resulted in a reduction of the amplitude difference of synaptic barrages simultaneously recorded in the apical dendrite and soma (Figure 3D), suggesting a greater electrical coupling between these two regions.

**Figure 3.**
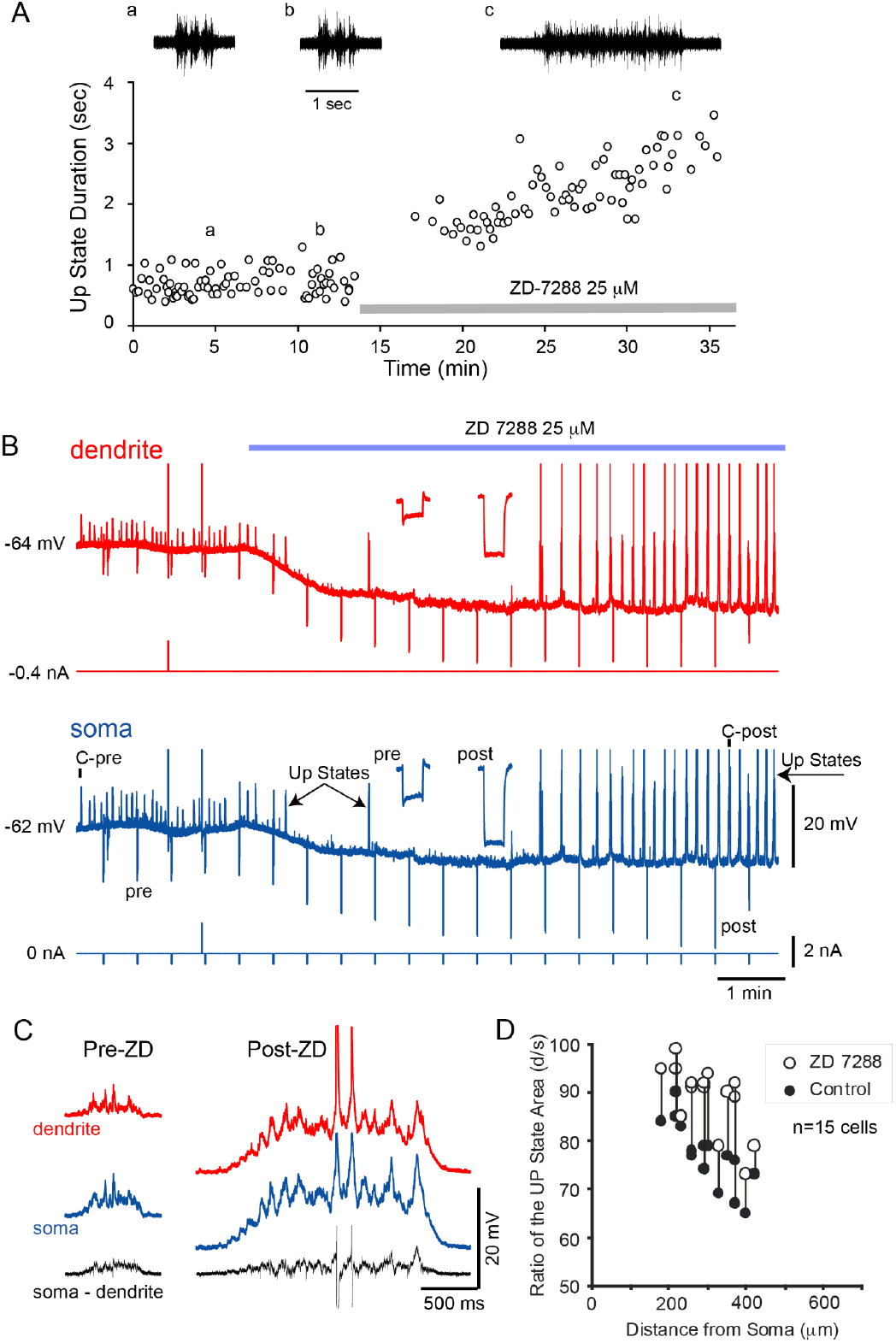
Application of the h-current blocker ZD7288 results in a progressive hyperpolarization of layer 5 pyramidal neurons and a large enhancement of Up-state synaptic barrages. A. Bath application of the h-current blocker ZD7288 (25 μM) results initially in a cessation of Up states, followed by their reappearance and prolongation in a multiple unit recording in layer 5 (submerged chamber). B. Simultaneous dendritic and somatic recording from a layer 5 pyramidal cell during the bath application of ZD7288. Note the pronounced hyperpolarization of both compartments. The Up states are initially blocked but return with a substantially increased amplitude and duration. The sag generated in response to a somatic hyperpolarizing current pulse is blocked following ZD7288. Note also the large increase in apparent input resistance. C. Example Up states recorded from the soma and apical dendrite before and after bath application of ZD7288. The bottom trace is the difference between the two compartments. D. Ratio of the Up state area recorded in the apical dendrite and soma before and after the bath application of ZD7288. Note that in every case, the reduction of the h-current brings the ratio closer to 100 percent.

Cesium is another blocker of the h-current. However, bath applications of Cs_^+^_ (1-2 mM) invariably resulted either in spreading depression (interface chamber; n=5) or marked increases in the rate of recurrence of the slow oscillation (submerged chamber; n=10). These responses were unrelated to the ability of Cs_^+^_ to block the h-current, since prior complete block of I_h_ with bath application of ZD7288 (25 μM) did not alter the Cs_^+^_-induced spreading depression or increased rate of Up state occurrence. These non-I_h_ effects of Cs_+_ therefore precluded its use in our experiments. The increase duration of the Up state following ZD7288 application was not merely a result of increased excitability, since increasing [K_+_]_o_ from 2.5 to 7 mM resulted in a progressive increase in the rate of the slow oscillation from one Up state every 10.6 (+/-7.4) to one every 0.6 (+/- 0.2) seconds. This increase in rate was associated with a decrease in the Up state duration from 0.41 (+/- 0.15) to 0.29 (+/- 0.14) seconds (n=4).

In addition to increasing the duration of the Up state, application of ZD7288 also increased the intensity of discharge of single pyramidal cells during the Up state, from an average firing rate of 1.8 +/- 1.7 Hz pre-ZD7288 to 9.2 +/- 4.9 Hz post ZD7288 (n=9, whole cell recordings). This increase in action potential activity occurred despite the hyperpolarization of the neurons, owing to the fact that the bath application of ZD7288 resulted in a marked increase in the amplitude of the PSP barrages arriving in both the somatic and dendritic recording sites (Figure 3B,C). In the somatic recordings, the PSP barrages during the Up state increased in amplitude, from an average peak of 7.6 +/- 2.7 mV to 20.7 +/- 7.2 mV (n=21; see Figure 3C). This increase in amplitude was larger than the hyperpolarization induced by ZD7288, indicating that the enhancement of the Up-state PSP barrage was not simply due to the increase in driving force. Indeed, returning the membrane potential of the recorded cell to pre-ZD7288 levels with somatic current injection revealed the PSP barrage to still be strongly enhanced in amplitude in comparison to pre-ZD7288 values (not shown; n=3). The greatly increased rapidity of the effects of ZD7288 in the submerged versus interface chamber presumably results from the increased ability of this agent to permeate the tissue in submerged slices.

In addition to hyperpolarizing the membrane potential, bath application of ZD7288 also resulted in a marked increase in apparent input resistance (from 50.7 +/- 9.6 to 68.0 +/- 13.1 MΩ, n=5, interface; soma: 18.6 +/- 7.5 to 36 +/- 10.8; dendrite: 23.1 +/-

10.1 to 46.2 +/- 17.0; n=10, submerged) as measured by small hyperpolarizing current pulses (Figures 3B, 4). This increase in apparent input resistance was present even following depolarization of the somatic compartment back to pre-ZD7288 levels (n=4; not shown). In conjunction with this increase in apparent input resistance, the depolarizing sag associated with the intracellular injection of hyperpolarizing current pulses, which is generated by the activation of the h-current (Solomon and Nerbonne, 1993; Solomon et al., 1993), was progressively blocked, for both dendritic and somatic current injections (Figures 3B, 4). Coincident with these effects, the bath application of ZD7288 also resulted in a prolongation of the membrane time constant, both as measured with the somatic recording electrode (from 9.9 +/- 2.0 to 21.1 +/- 9.8 msec) and dendritic recording electrode (6.0 +/- 2.0 to 21.1 +/- 9.2 msec; n=10).

The intracellular injection of depolarizing current pulses into either the somatic or apical dendritic compartments revealed that the bath application of ZD7288 resulted in a reduction in the threshold current needed to initiate an action potential as well as a marked increase in the number of action potentials generated in response to supra-threshold current pulses, even without compensation for the hyperpolarizing effect of h-current block (Figure 4B, E; n=9). This effect could be so pronounced that either the somatic or dendritic injection of a current pulse that was just threshold for the generation of one action potential before the application of ZD7288 would activate a train of action potentials following the application of this drug (Figure 4B).

**Figure 4.**
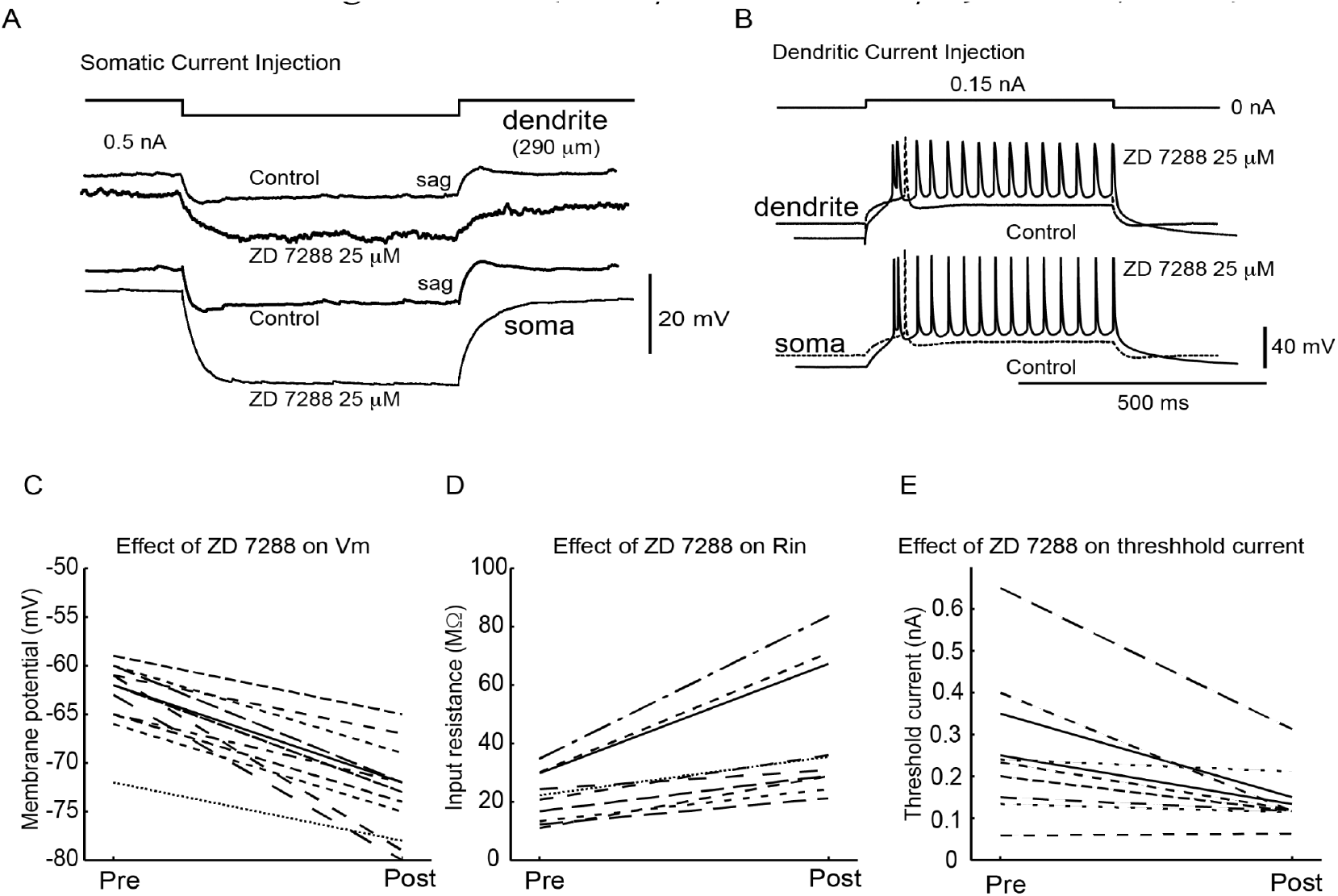
Reduction of the h-current hyperpolarizes membrane potential, increases the apparent input resistance, and increases neuronal responsiveness. A. Hyperpolarizing response in the apical dendrite and soma in response to the injection of a hyperpolarizing current pulse into the dendrite. Bath application of ZD7288 results in an abolition of the depolarizing sag as well a large increase in the apparent input resistance. B. Response of the neuron to the intra-dendritic injection of a depolarizing 0.15 nA current before and after the bath application of ZD7288. The reduction of the h-current results in a hyperpolarization of the cell, yet the neuron’s responsiveness is greatly enhanced such that instead of one spike (dashed line), the dendritic current pulse generates a prolong train of spikes (solid line). C, D, E. Effect of ZD7288 on the resting membrane potential, apparent input resistance, and threshold current needed to initiate an action potential. All cells recorded in the submerged chamber.

Examination of the f-I relationship for somatic current injection revealed that bath application of ZD7288 resulted in an increase in average slope from 25.6 (+/-5.6) to 40.2 (+/- 8.5) Hz/nA (p<0.05, n=9; see Supplement – 3; firing rate is the average rate during a 500 msec pulse).

These results indicate that the ability of ZD7288 to enhance the amplitude and duration of the PSP barrages arriving in cortical pyramidal cells resulted at least in part from the increased apparent input resistance and lengthened membrane time constant of these neurons. However, since single pyramidal neurons increased their firing rate (averaged over the entire period of the Up state) following the application of ZD7288, we would expect that average network activity, as represented by synaptic barrages, would also be enhanced. To examine whether or not the synaptic currents underlying the barrages were enhanced in some manner following the block of the h-current, we performed somatic whole cell voltage clamp recordings with electrodes containing QX-314 (1 mM) to block active Na_^+^_ and voltage clamped the recorded cell to membrane potentials varying from near the reversal potential for IPSCs (–75 mV) to the reversal potential for EPSCs (0 mV) (Shu et al., 2003a). The combined influence of voltage clamp as well as the ability of QX-314 to reduce the h-current(Perkins and Wong, 1995) should minimize the effects of changes in properties of the recorded cell by ZD7288 on the observed synaptic currents. Under these conditions, the bath application of ZD7288 resulted in a marked enhancement of the synaptic currents recorded both at –75 and 0 mV, when comparing the same time period of the Up state before and after ZD7288 application (Figure 5). Excitatory PSC dominated traces recorded at –75 mV averaged – 0.14 +/- 0.13 nA for the first 1000 msec of the Up state in control, and this was increased to –0.27 +/- 0.1 nA for the same time period following the bath application of ZD7288. Similarly, ZD7288 enhanced IPSC barrages (recorded at 0 mV) from an average of 0.25+/- 0.16 to 0.65 +/- 0.27 nA (Figure 5; n=5). Closer examination of the traces reveals that the enhancing effect of ZD7288 was not due to an increase in the peak amplitude of the EPSC or IPSC barrages, but rather a decrease in the rate at which these PSC barrages decreased in amplitude during the Up state. These results are consistent with an enhancing effect of ZD7288 on cortical recurrent network activity

**Figure 5.**
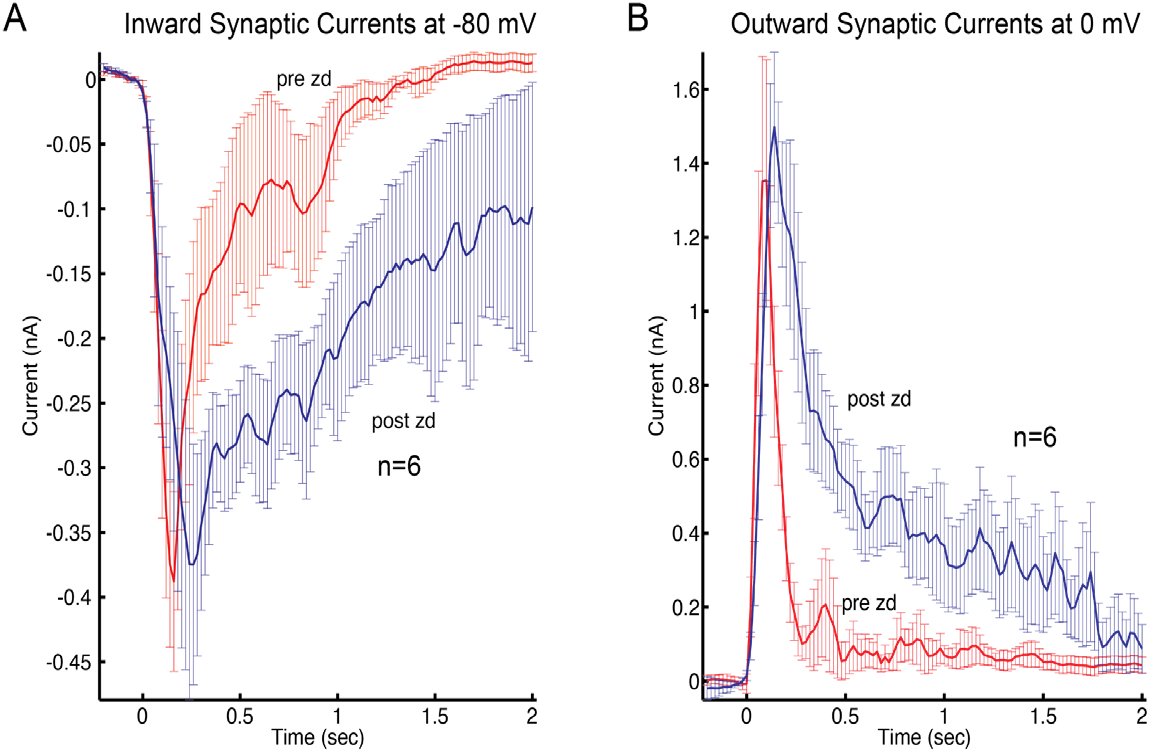
Bath application of ZD7288 enhances the maintenance of synaptic activity in cortical networks. A. Inward currents recorded at -80 mV in layer 5 pyramidal cells (n=6) during Up states before and after bath application of ZD7288 (25 μM). At this membrane potential, the synaptic currents should be dominated by EPSCs. B. Synaptic currents recorded while the cells were voltage clamped near 0 mV. At this membrane potential, the synaptic currents will be dominated by IPSCs. Note that both EPSCs and IPSCs are more prolonged following the bath application of ZD7288. Error bars are standard error of the mean.

### Action potentials back-propagate into the apical dendrite

Prior to the application of ZD7288, when the Up state resulted in the initiation of action potentials, these always occurred first in the somatic compartment and propagated at a rate of 0.66 +/- 0.23 m/sec into the apical dendrite (Figure 6A,C; 332 spikes in 16 cells). Similarly, the intracellular injection of a depolarizing current pulse into the soma also caused action potentials in the somatic compartment that back-propagated into the apical dendrite, even though local depolarization of the apical dendrite with current injection could initiate action potentials locally that then propagated into the somatic compartment (not shown; (Berger et al., 2001)). Following the block of the h-current, Up-state initiated action potentials were still nearly always generated first in the somatic/initial segment/axon compartment, and back-propagated into the apical dendrite, although occasionally spikes were initiated in the apical dendrite and then propagated into the somatic region (Figure 6B,D). The application of ZD7288 had a significant effect on action potential shape in the dendritic compartment and promoted the occurrence of burst-discharges (Figure 6; see Supplement-4).

**Figure 6.**
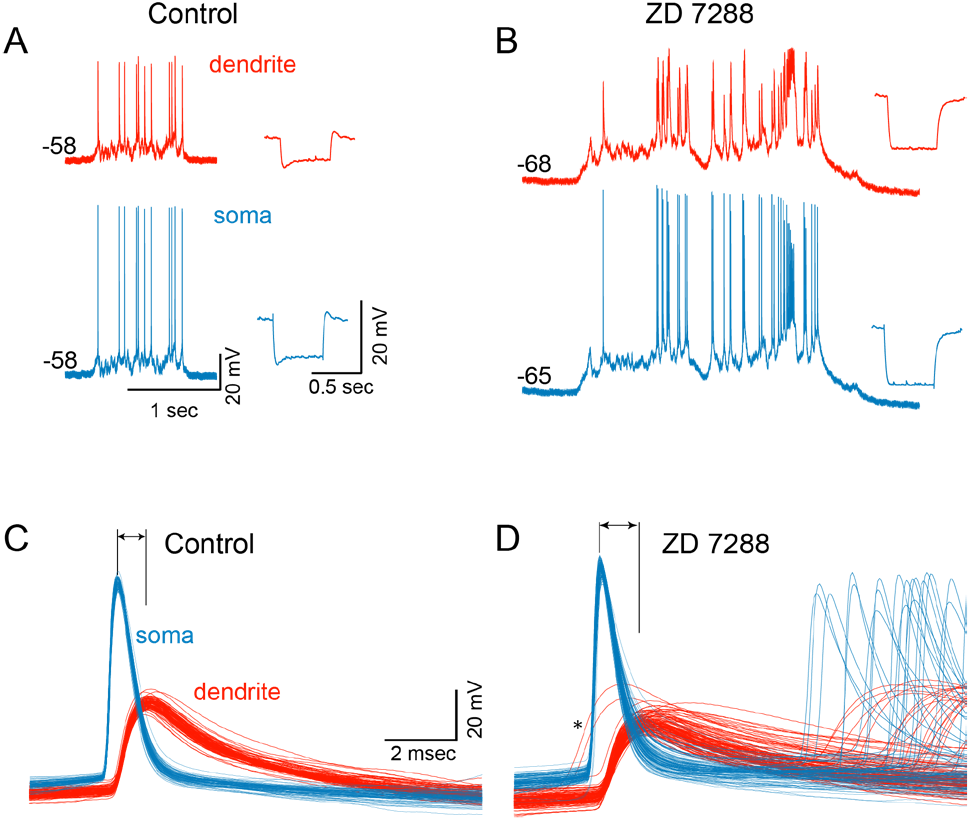
Action potentials backpropagate from the somatic to apical dendritic segments during Up states before and after block of the h-current. A. Example of a simultaneous recording of an Up state in the soma and apical dendrite of a cell before application of ZD7288. Expansion of the action potentials reveals that they all occur in the somatic compartment prior to the apical dendrite (C). B. Same recording as A, except following the block of the h-current with the bath application of ZD7288. As before the application of ZD7288, almost all Up state action potentials are generated in the somatic prior to the apical dendritic compartments (D). Two action potentials appear to occur in the dendrite first, followed by spiking in the soma (asterisk). C, D. Overlay of action potentials during Up states. Action potentials were chosen such that at least 30 msec had passed since the last action potential. The application of ZD7288 appears to broaden dendritic action potentials as well as increase the incidence of burst discharges.

Although the bath application of ZD7288 resulted in only a small, but insignificant, prolongation of somatic action potentials recorded in either the interface (1.7 +/- 0.15 to 1.8 +/- 0.17 msec) and submerged (1.3 +/- 0.13 to 1.4 +/- 0.23 msec; n=4) chambers, the application of this h-current blocking agent resulted in a significant prolongation of action potentials recorded from the apical dendrite from 1.93 +/- 0.16 msec to 2.62 +/- 0.3 msec (P< 0.05; n=5) (only action potentials that occurred at least 30 ms after the preceding action potential were included for analysis, to eliminate action potentials generated after the start of a burst). In submerged chamber recordings, the application of ZD7288 typically resulted in the appearance of burst discharges in regular spiking neurons (n=17/19 cells), presumably owing to the prolongation of dendritic action potentials (see Supplement 4; this effect was not observed in neurons recorded in the interface chamber; n=5) (see also (Berger et al., 2001)).

The block of the h-current with ZD7288 did not affect action potential threshold, either that associated with action potentials generated by spontaneous synaptic barrages of the Up state (−45.2 +/- 6.8 mV pre-ZD7288, -44.3 +/- 11.7 mV post; n=4; submerged chamber), or with the intracellular injection of current ramps (−54.5+/-3.2 mV pre, - 56.7+/- 4.7 mV post-ZD7288; n=5; interface chamber; see Supplement 5)

### Reduction of h-current enhances dendrosomatic transfer in a frequency dependent fashion

Previous investigations have demonstrated that a reduction of the h-current results in a slowing of the membrane time constant and consequently an increase in temporal summation in the apical dendrites of pyramidal cells (Magee, 1998, 1999; Williams and Stuart, 2000; Berger et al., 2001, 2003; Nolan et al., 2004; Harnett et al., 2015). In addition, these studies have demonstrated that the h-current potently controls the ability of single EPSPs or trains of EPSP-like events to depolarize and initiate action potentials in the somatic compartment. Here we explored the effects of h-current blockade on dendrosomatic and somatodendritic communication by determining the conductance to voltage transfer function between the apical dendritic and somatic compartments. To calculate this transfer function, we injected into either the somatic or apical dendritic recording electrode in vivo-like excitatory conductances that contained a broad range of frequencies (see Methods) with the dynamic clamp technique (Destexhe et al., 2001; Dorval et al., 2001; Shu et al., 2003a; Zerlaut et al., 2016). The resulting changes in membrane potential at both recording sites were then measured, and the resulting transfer functions calculated (Figure 7; see Methods).

**Figure 7.**
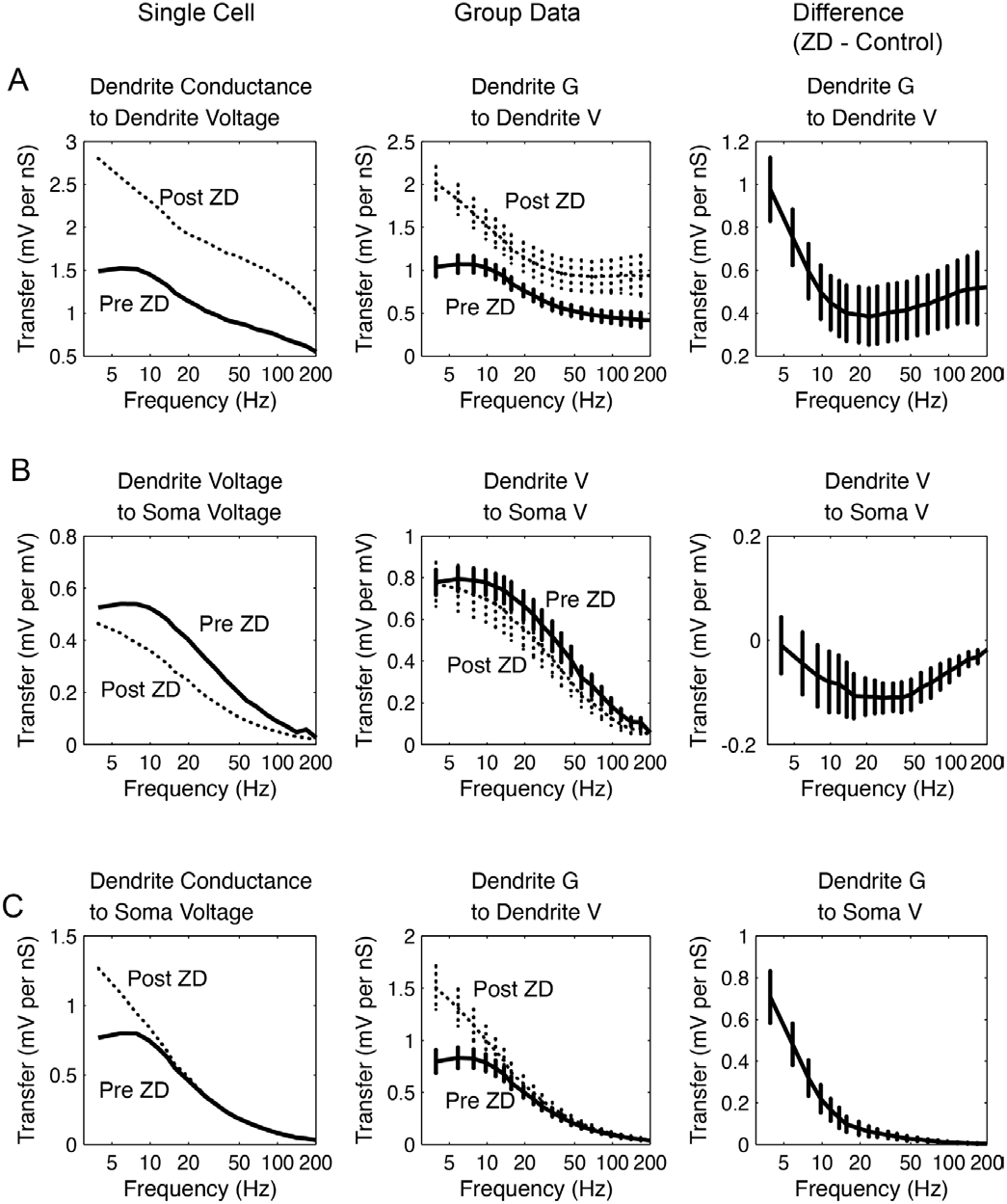
Reduction of the h-current enhances the transfer of dendritic conductances to somatic voltage in a frequency dependent manner. A. Single cell and group data showing that ZD7288 enhances the induction of voltage deviations by locally injected conductances (injected into the apical dendrite) especially at frequencies below approximately 15 Hz. The plot on the right is the difference between the pre- and post-ZD7288 group data traces. B. Single cell and group data demonstrating that ZD7288 reduces the transfer of dendritic voltages to somatic voltages, with a peak decrease at 15-50 Hz. C. Single cell and group data demonstrating that the transfer of dendritic conductance to somatic voltage is enhanced by ZD, particularly for frequencies below approximately 20 Hz. All data obtained with dual apical dendritic and somatic patch clamp recordings and injection of broad band conductance noise into the dendritic recording site. In the example cell, the apical dendritic recording site was 350 μm from the soma. Error bars are standard error of the mean.

First, we calculated the frequency-dependent transfer from local conductance to local voltage in both the apical dendrite and somatic recording electrodes, before and after application of ZD7288 (25 μM in bath; Figure 7A; n=6). We found that block of the h-current enhanced the ability of dendritically injected conductances to drive changes in membrane potential across a broad spectrum of frequencies (5-200 Hz), although the enhancement was largest for lower frequencies (Figure 7A). The effect of ZD7288 on the transfer of voltage between the dendritic and somatic compartments was also frequency dependent. Reduction of the h-current resulted in a decrease in transfer of voltage frequencies that peaked at about 20 Hz, with lesser effects at both lower and higher frequencies (Figure 7B). Note that the magnitude of this effect was less than the effect of ZD7288 on the transfer of local dendritic conductance to voltage. This broadband enhancement of local conductance to voltage transfer, together with a weaker frequency dependent attenuation of voltage transfer from dendrite to soma, resulted in a frequency dependent enhancement in transfer of dendritic conductance to somatic voltage (Figure 7C). Thus the block of the h-current with ZD7288 resulted in a marked enhancement of power transfer that was most marked below approximately 20 Hz (P<0.01; n=6), with a smaller enhancement of frequencies between 20 and 140 Hz (Figure 7C).

The Up state of the slow oscillation is generated through recurrent excitation in local cortical networks (Steriade et al., 2001; Shu et al., 2003a, 2003b). The ability of ZD7288 to strongly enhance the duration of the Up state, and the amplitude of PSP barrages arriving at the soma and apical dendritic recording sites, suggests that the h-current may play an important role in the communication of ongoing synaptic activity throughout the cell. To test this hypothesis, we examined the effect of ZD7288 on Up-like states that were injected into either the dendritic or somatic recording electrode using the dynamic clamp technique. We found that block of the h-current with bath application of ZD7288 (n=5) resulted in a marked enhancement of the transmission of Up-like states from the apical dendrite (recorded at an average distance of 217 μm) into the somatic compartment, such that a greater number of action potentials were generated in response to these dendritic conductance injections (Figure 8). Similarly, bath application of ZD7288 also greatly enhanced the ability of somatically injected artificial Up-states to initiate action potentials (not shown; n=4), even in the presence of the pronounced hyperpolarization resulting from ZD7288 application.

**Figure 8.**
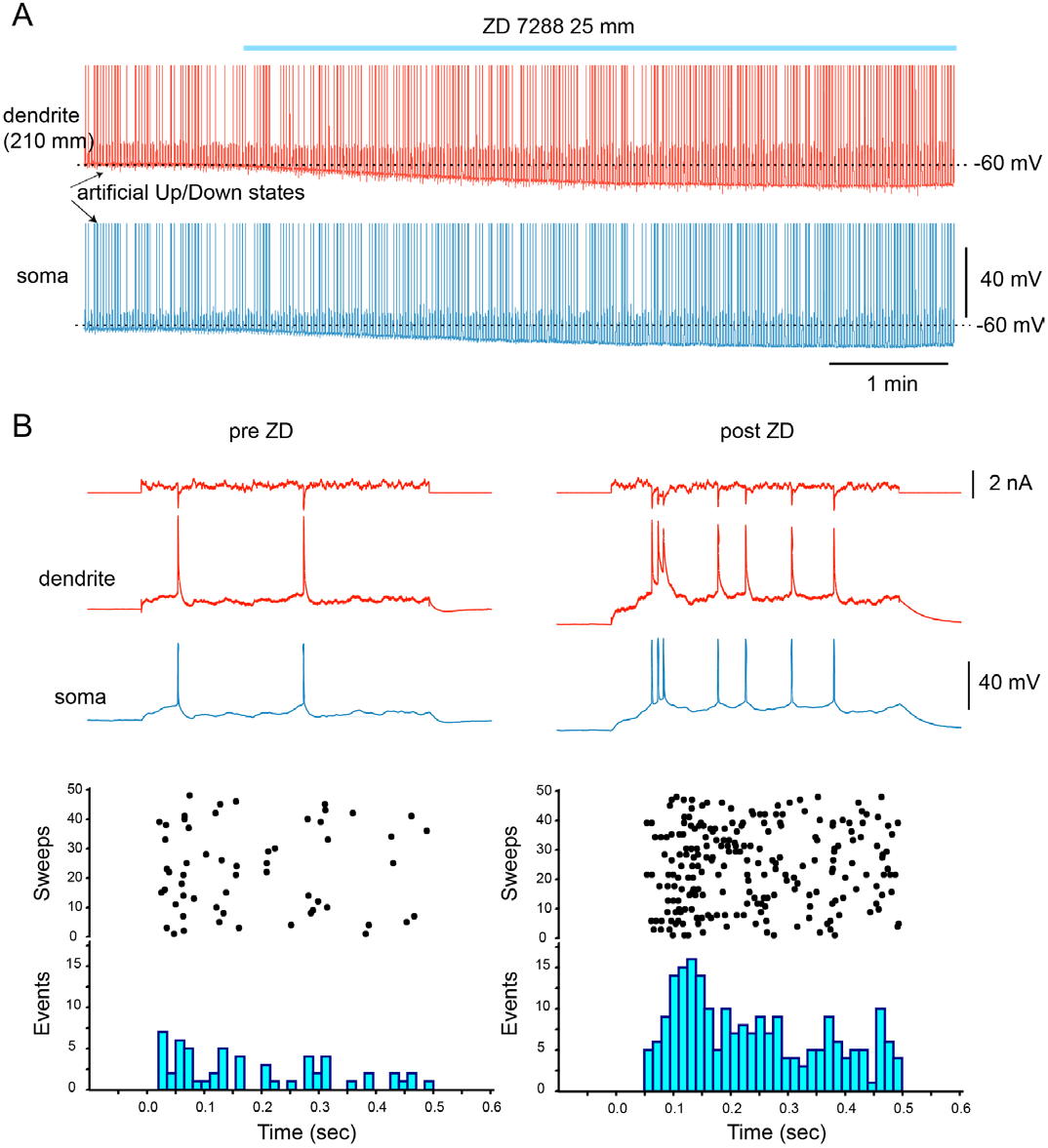
Reduction of the h-current results in an enhanced response to the dendritic injection of Up-state like currents. A. Artificial Up-like states were generated through the injection of a balanced excitatory and inhibitory synaptic-like conductance into the apical dendrite using a dynamic clamp system. Bath application of ZD7288 results in a hyperpolarization of the membrane potential, but an increased ability of these artificial dendritic Up states to initiate action potentials. B. Examples of the response to the injection of artificial Up states before and after the application of ZD7288. Histograms at the bottom illustrate that the action potential response to the artificial dendritic Up state is enhanced following the block of the h-current.

## Discussion

A characteristic anatomical feature of the neocortex is the presence of massive recurrent networks, both locally, and more globally. Although the functional operation of these networks has long been proposed to mediate important cortical functions (Pinto et al., 2022; Lorente De No, 1938; Hebb, 1949; Wang, 2003; Major and Tank, 2004; Funahashi and Procyk, 2022), the cellular details of the operation of cortical recurrent networks have only recently begun to be explored. One relatively easily studied example of recurrent network neocortical activity is the generation of Up and Down states during slow wave sleep, anesthesia, and in vitro (Steriade et al., 1993, 2001; Cowan and Wilson, 1994; Shu et al., 2003a). Here we have developed a submerged cortical in vitro slice preparation that allowed us to examine the influence of the h-current on the generation of the recurrent network activity of the Up state, and the potential role of dendrosomatic communication in this effect.

The h-current is a time-dependent hyperpolarization-activated Na_^+^_ /K_^+^_ current that contributes strongly to the resting membrane potential, input resistance, and pacemaker properties of neurons (reviewed in (Pape, 1996; Lüthi and McCormick, 1998; He et al., 2014)). Immunocytochemical staining for the four subunits of the h-current (HCN1-4) in the rat neocortex have revealed the strong presence of both HCN1 and 2, particularly in the apical dendrites of layer 5 pyramidal neurons, where the density of these subunits, and of the h-current, steadily increases towards layer 1 (Lörincz et al., 2002; Notomi and Shigemoto, 2004). The block of the h-current results in significant changes in the integrative properties of apical dendrites, prolonging and increasing the summation of excitatory postsynaptic potentials (Schwindt and Crill, 1997; Magee, 1998, 1999; Stuart and Spruston, 1998; Williams and Stuart, 2000, 2003; Berger et al., 2001, 2003; Nolan et al., 2004; Oviedo and Reyes, 2005; Sheets et al., 2011). Thus, the h-current controls the integrative properties of the apical dendrites of at least some types of layer 5 pyramidal neurons. Here we demonstrate that the block of the h-current with ZD7288 results in a pronounced prolongation of the recurrent network activity that underlies the Up state. This enhancement of recurrent network activity presumably results from the enhanced ability of somatic and dendritic depolarizations to initiate action potentials, despite the hyperpolarization associated with reduction of the h-current.

Although at high concentrations (> 50 μM) ZD7288 can have effects in addition to those of block of the h-current, such as the antagonism of glutamatergic receptors and subsequent reduction of EPSPs (Chevaleyre and Castillo, 2002; Chen, 2004), our results are consistent with enhancement of the Up state through reduction of the h-current.

The bath application of ZD7288 was effective at concentrations as low as 1 μM and resulted in a membrane hyperpolarization, increase in apparent input resistance, and block of the depolarizing sag activated by hyperpolarizing current pulses. These effects are all consistent with those expected from reduction of the h-current, and occurred with a time course that was similar to the period in which the Up state duration was lengthened. Reduction of glutamatergic transmission, on the other hand, reduces or blocks the generation of Up states (Sanchez-Vives and McCormick, 2000), indicating that the enhancing effect of ZD7288 on Up states is not mediated through this mechanism.

Up states are generated through barrages of synaptic activity in which recurrent excitation and inhibition are proportional and balanced (Shu et al., 2003b). This proportionality results in a reversal potential that is very steady even during large changes in total synaptic conductance. The presence of strong barrages of excitatory and inhibitory postsynaptic potentials in every cortical neuron recorded (Steriade et al., 1993, 2001; Hasenstaub et al., 2005), the significant numbers of cortical cells that discharge with the generation of each Up state, as indicated by extracellular multiple unit recordings (Hasenstaub et al., 2005), and the complete block of this activity by antagonists of excitatory synaptic transmission (Sanchez-Vives and McCormick, 2000), indicate that recurrent synaptic activity is the major component in the generation of this recurring, persistent activity. This being said, it is also clear that intrinsic cellular properties necessarily contribute to recurrent cortical activity. In particular, the ease with which synaptic barrages can initiate action potentials in the postsynaptic cells is critical. The h-current controls this synaptic barrage-action potential coupling through at least three mechanisms: determination of the membrane potential, of input resistance, and of integrative time constant. We have found that although reductions in the h-current result in hyperpolarization of the membrane potential, the threshold somatic or apical dendritic current for action potential initiation is reduced, presumably due to increased efficiency of coupling between injected current and spike initiation.

Since spikes are typically initiated in the axon hillock/somatic compartment and then back-propagate into the dendrites (Stuart et al., 1997a, 1997b; Larkum et al., 2001; Foust et al., 2010; Popovic et al., 2011), this implies that reduction of the h-current increases the efficiency with which dendritic or somatic depolarizations activate the axon hillock. This increased ability of injected currents to initiate action potentials results, presumably, from the concomitant increase in input resistance and increased ability of current to affect a larger region of the cell (e.g. lengthened membrane space constant). Although this is particularly true for synaptic inputs arriving in the distal apical dendrite (owing to the high concentration of h-channels in this location) (Lörincz et al., 2002; Notomi and Shigemoto, 2004; Harnett et al., 2015), it is also valid for other parts of the cell, since h-channel activity in the apical dendrite affects the integrative properties of the entire neuron. Reduction of the h-current results in a pronounced, but frequency dependent, enhancement of conductance to voltage transfer. The transfer of lower frequencies is much more strongly enhanced by block of the h-current than the transfer of higher frequencies, owing to the lengthening of the membrane time constant, which alters the temporal and spatial filtering properties of the neuron. This alteration preferentially enhances prolonged depolarizations, such as those of the Up state, since most of their power is in low frequencies (<1 Hz).

Computational models and in vitro data suggest that Up states are generated through a balance of recurrent excitation/inhibition offset by: 1) refractory mechanisms such as the buildup of outward currents from synaptic and action potential activity (Sanchez-Vives and McCormick, 2000; Compte et al., 2003); 2) increases in the activation of inhibitory pathways (Brunel and Wang, 2001; Shu et al., 2003b); 3) depression of excitatory synapses (Bazhenov et al., 2002); or 4) increased synchrony of action potential activity (Gutkin et al., 2001). Once a critical juncture is obtained in which the inward currents that maintain the Up state are no longer sufficient to offset the combined “inhibitory” influences of synaptic inhibition combined with the refractory mechanisms, then the Up state suddenly fails and the network rapidly transitions into the Down state. We propose that reduction of the h-current enhances the ability of the network to maintain recurrent excitatory activity through an increase in the ability of depolarizing synaptic barrages to initiate action potentials, in at least layer 5 cortical pyramidal cells (see also (Sheets et al., 2011)). The reduction in h-current would be particularly enhancing to the effectiveness of excitatory PSPs, since, on average, these are more distal to the soma/axon initial segment than are inhibitory inputs, and therefore more prone to reduction in amplitude owing to the shunting effects of the h-current. We have shown previously that layer 5 is a key site for the generation of Up states in vitro, since this activity occurs earliest and is strongest in layer 5, and surgical separation of the infragranular layers from the supragranular layers leaves Up and Down states intact only in the forme r(Sanchez-Vives and McCormick, 2000). We envision that this increased efficiency with which depolarization influences activation of action potentials results in a marked prolongation of the duration of Up states, since it will take longer for the refractory mechanisms, regardless of what they are, to build up to the point where they effectively offset the increased efficacy of excitation. Another interesting possibility is that reductions of the h-current result in the reduced propensity for neuronal networks to generate synchronized activity (Acker et al., 2003), thereby resulting in an increase in duration of both the Up and Down states, since state transitions may correspond to critical levels and patterns of neuronal synchrony (Gutkin et al., 2001; Shu et al., 2003b).

Interestingly, the voltage dependence of the h-current is directly modulated by cAMP, although to varying degrees depending on subunit composition and post-translational modifications (Ludwig et al., 1998, 1999; Santoro et al., 1998; Ishii et al., 1999; Seifert et al., 1999; Ulens and Tytgat, 2001; Chen, 2004). A variety of neurotransmitters can modulate cAMP levels and the h-current in cortical neurons (Bickmeyer et al., 2002; Poolos et al., 2002; Schweitzer et al., 2003; Barth et al., 2008; Heys and Hasselmo, 2012; Grzelka et al., 2017; Labarrera et al., 2018). We hypothesize that the ability of cortical networks to generate recurrent network activity is potently controlled through neurotransmitter modulation of the h-current via cAMP mechanisms. The ability of cortical networks to rapidly form and break functional microcircuits through dynamically interacting recurrent networks is believed to be a key mechanism by which the neocortex operates, and may provide the neural substrate for working memory (see (Durstewitz et al., 2000; Wang, 2001; Major and Tank, 2004)), attention (Chance et al., 2002; Shu et al., 2003a; Haider and McCormick, 2009), context, and behaviorally relevant sensory-motor coupling (Andersen and Mountcastle, 1983; Andersen et al., 1985). Modulation of the h-current may, therefore, have a strong influence on the formation and dissolution of coactive neuronal networks (e.g. ensembles) during behavior.

## Acknowledgements

Supported by the NIH R35NS097287, RF1NS118461, R37NS026143

