## Supplemental Figures for "The h-current controls cortical recurrent network activity through modulation of dendrosomatic communication"

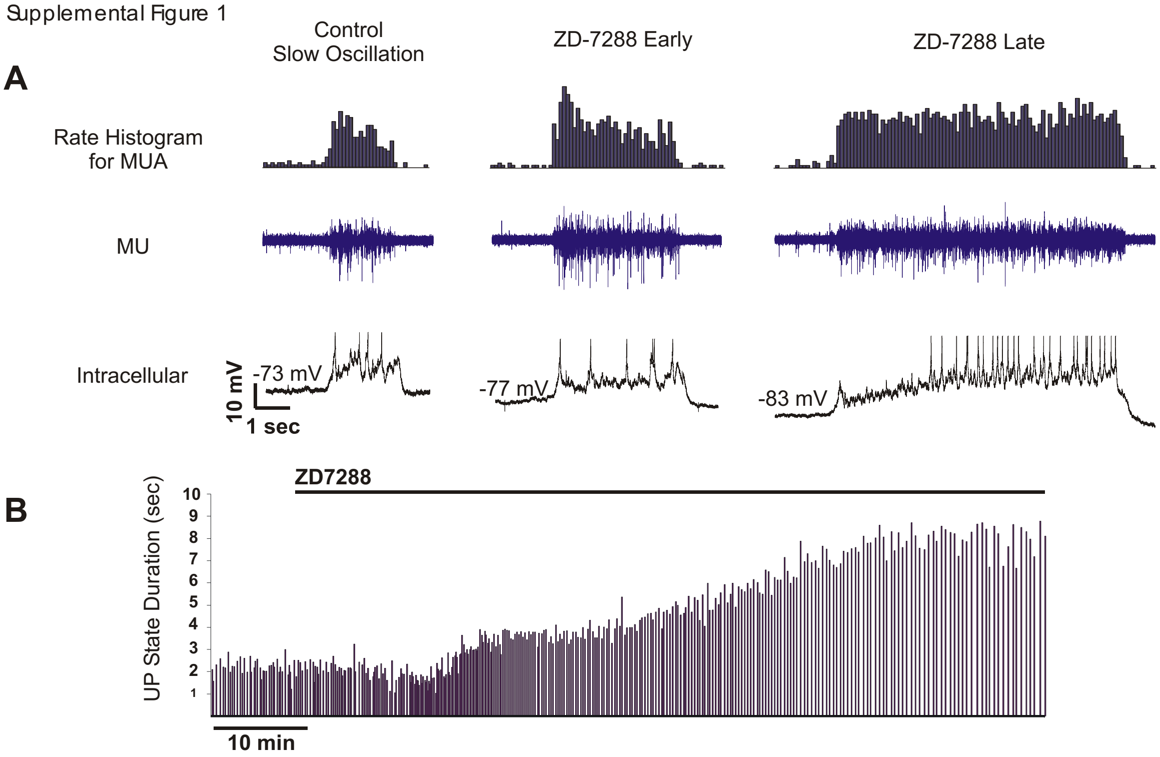


Figure 1. Bath application of ZD-7288 results in the gradual prolongation of Up states in prefrontal cortical slices in the interface chamber. A. Simultaneous extracellular multiple unit (MU) and intracellular recording from layer 5 before and during wash in the h-channel blocker ZD7288 (100 µM). Note the marked prolongation of Up states in both the extracellular and intracellular recordings and the increased firing rate of the intracellularly recorded neuron. B. Plot of Up state duration versus time before and after bath application of ZD7288. Similar effects were found with 20 and 50 µM bath concentrations of ZD-7288. The effects of ZD7288 occurred only very slowly in the interface, compared to the submerged, recording chambers, presumably owing to slow and incomplete drug penetration of the tissue in the interface chamber.

Supplement Figure 2

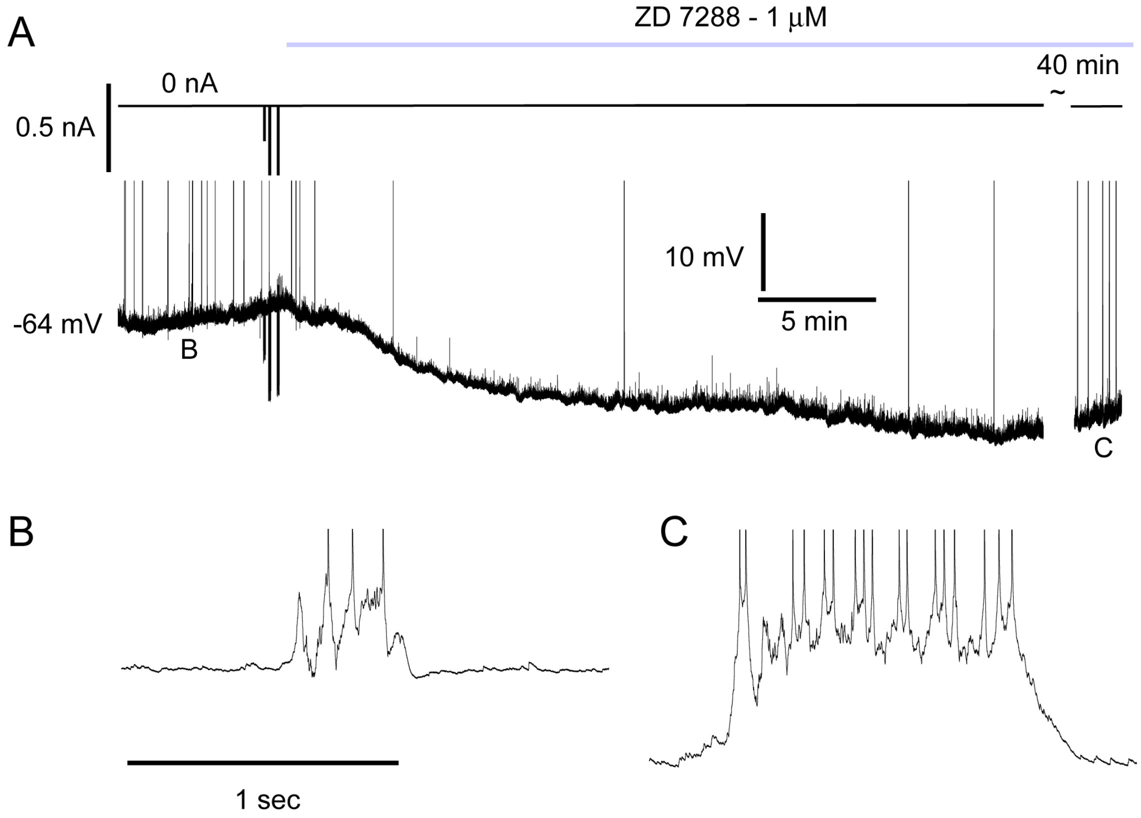


Figure 2. Application of low doses of ZD7288 also hyperpolarize layer 5 pyramidal neurons and increase the duration of Up states. A. Intracellular recording from a layer 5 pyramidal cell in a submerged chamber. Bath application of 1 µM ZD7288 results in a hyperpolarization of the neuron and an initial decrease in the rate of UP state generation. The Up state generation returned to normal, although the duration and amplitude of each Up state was greatly increased, as with higher doses of ZD7288. B and C. Expansion of Up states before and after wash in of 1 µM ZD7288.

Supplement Figure 3

Effects of ZD-7288 on the f-I relationship of Cortical Pyramidal Cells

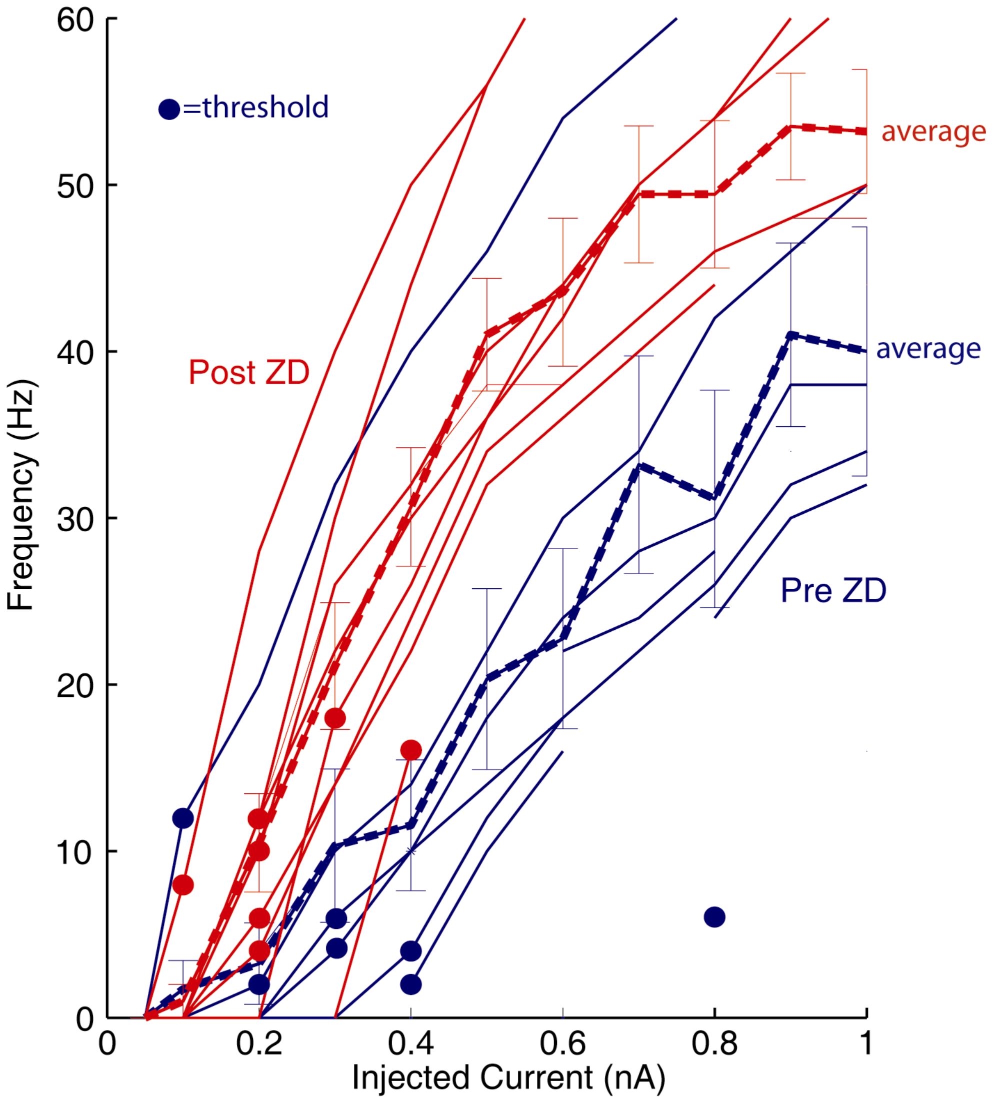


Figure 3. Application of the h-channel blocker ZD7288 results in an increase slope of the f-I relationship of layer 5 cortical pyramidal cells. Blue traces are before, while red traces are after, bath application of ZD7288 (25 μM). Thick dashed lines are the average of the population (n=9), and error bars are expressed as standard error of the mean. Filled dots represent the first current pulse to evoke action potentials for each cell. Frequency of firing is the average rate over the 500 msec current pulse. Changes in membrane potential were not compensated for with the intracellular injection of DC.

Supplemental Figure 4

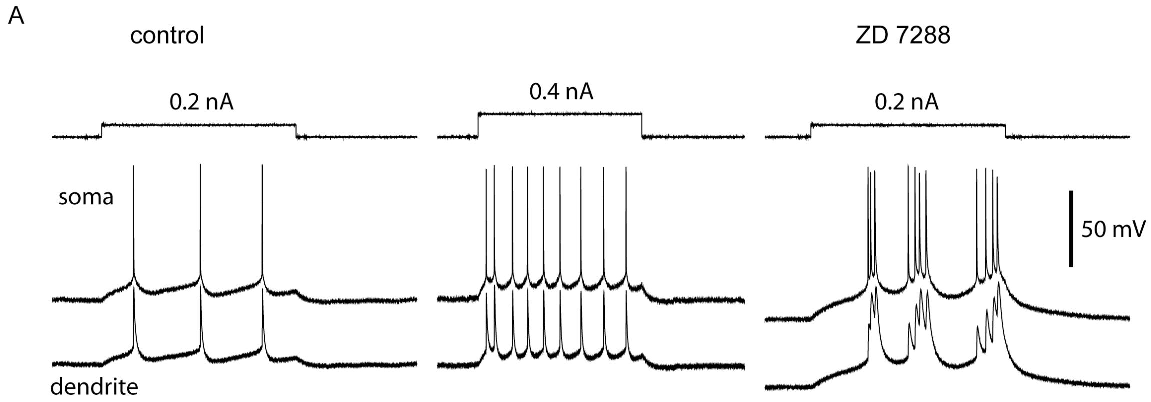


Figure 4. Bath application of ZD7288 increases the incidence of burst firing in layer 5 pyramidal neurons. A. Response of a layer 5 neuron, as recorded both in the soma and apical dendrite (350 μm from the soma), to the intracellular injection of 0.2 and 0.4 nA depolarizing current pulse injected into the soma. After the block of the h-current with ZD7288 (25 μM), current pulses result in the generation of bursts of action potentials in both the somatic and dendritic recording electrodes. This effect was observed in 17/19 neurons recorded in the submerged chamber.

Supplemental Figure 5

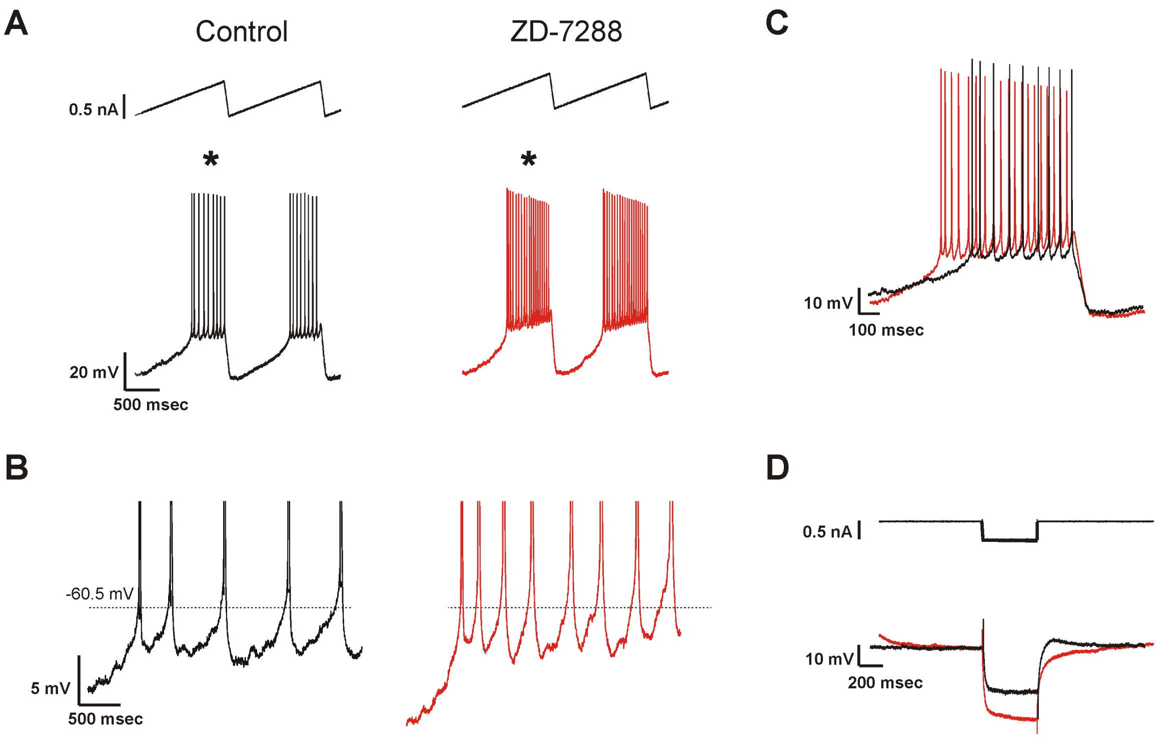


Figure 5. Block of the h-current with bath application of ZD7288 results in an increase in responsiveness, but does not alter spike threshold. A. Response of a layer 5 regular spiking neuron to the intracellular injection of a depolarizing ramp before and after bath application of ZD-7288 in an interface chamber. B. Expansion of the traces indicated by an asterisk. The dashed line represents action potential threshold for the first action potential. C. Overlap of the response to the indicated depolarizing ramps in A. The bath application of ZD7288 results in a marked increase in responsiveness, but not a change in action potential threshold. D. Overlap of the response of the neuron to hyperpolarizing current pulses before and after the bath application of ZD7288. Although this neuron does not exhibit a prominent depolarizing sag upon hyperpolarization, there is a depolarizing after potential that is abolished by application of ZD7288.
